# ‘Formation and function of the meninges arachnoid barrier around the developing brain’

**DOI:** 10.1101/2022.06.10.495709

**Authors:** Julia Derk, Christina N. Como, Hannah E. Jones, Luke R. Joyce, Stephanie Bonney, Rebecca O’Rourke, Brad Pawlikowski, Kelly S. Doran, Julie A. Siegenthaler

## Abstract

Barriers at the level of the brain endothelium, choroid plexus, and meninges strictly regulate movement of molecules and cells into and out of the central nervous system (CNS). In contrast to the blood-brain barrier and choroid plexus epithelial barrier, developmental timing and function of the meningeal arachnoid barrier, a layer of epithelial-like cells connected by tight and adherens junctions, is largely unknown. To begin to address this, we mined our E14.5 mouse single cell transcriptomic (scRNA-seq) meningeal fibroblast data set and identified the repression of Wnt-β-catenin signaling as a key mechanism underlying the specification of epithelial-like arachnoid barrier cells from Collagen 1+ and Crabp2+ mesenchymal meningeal precursors. We show that elevating Wnt-β-catenin signaling in prenatal meningeal mesenchymal cells prevented the development of arachnoid barrier cells. In the absence of dorsal arachnoid barrier cells, the prenatal meninges and brain are penetrable to biocytin-TMR and *Streptococcus agalactiae* (Group B *Streptococcus*, GBS), the leading pathogen known to drive life-threatening neonatal meningitis. We show that a layer of Claudin 11 (tight junction) and E-cadherin (adherens junction) expressing arachnoid barrier cells appear around the mouse brain from E13-E15 and the emergence of a functional barrier by E17 coincides with junctional localization of Claudin 11. Postnatal growth of the arachnoid barrier is marked initially by proliferation and later re-organization of junctional domains. This work provides fundamental knowledge on development and prenatal function of a meningeal arachnoid barrier, and novel tools for future studies on regional functions of this CNS barrier in the meninges.

## Introduction

The meningeal arachnoid barrier is a component of the blood-cerebrospinal fluid barrier (B-CSFB), that along with barriers at the pial vasculature and choroid plexus, regulates the free movement of molecules and cells into and out of the CSF in the subarachnoid space and ventricles (Balin et al., 1986; Ichikawa and Itoh, 2011; Ichikawa et al., 2010; Rodriguez-Peralta, 1957; Roth et al., 2013). A mechanistic understanding of how the blood-brain barrier (BBB) and B-CSFB develop and function is important, as their disruptions contribute to the pathophysiology of brain injury, neurodegenerative disorders, neuroinflammatory diseases, cancer, and the aged brain (Alves de Lima et al., 2020; Brkic et al., 2015; Derk et al., 2021; Iliff et al., 2012; Liebner et al., 2018; Sweeney et al., 2019). The arachnoid barrier separates the fenestrated blood vessels of the dura from the immune privileged CSF and is composed of epithelial-like cells connected by tight and adherens junctions (Nabeshima et al., 1975; Polanco et al., 2021; Rascher and Wolburg, 1997; Uchida et al., 2019). Our prior work showed that arachnoid barrier cells have a shared mesenchymal origin with other meningeal fibroblasts at E12 and begins to express epithelial (E-cadherin) and tight junction (Claudin 11) proteins in the prenatal meninges with a discontinuous layer at E14 (DeSisto et al., 2020). Similar expression of Claudin 11 layer are also seen in the human and rat fetal brain (Brøchner et al., 2015). Here we define the developmental mechanisms that drive meningeal mesenchymal precursor cells to transition into epithelial-like arachnoid barrier cells with junctional properties around the pre-and postnatal CNS. We show that a downregulation of Wnt-β-catenin signaling is a key molecular mechanisms behind arachnoid barrier cell specification and that the barrier is functional as early as E17 to protect the CNS from invasion from peripheral molecules and foreign pathogens in late prenatal stages. These studies significantly advance our knowledge on CNS barrier development and generates novel tools and mechanistic insights that will empower future studies to uncover how the B-CSFB, and specifically the arachnoid barrier, contributes to CNS health and disease.

## Results

At E14 arachnoid barrier cell specification is underway but incomplete, as arachnoid barrier cells (defined by E-cadherin expression), are present in the ventral meninges but are not yet observed in the dorsal meninges (DeSisto et al., 2020). To compare the transcriptional signature of mature arachnoid barrier cells to those that are actively specifying, we took our previously generated E14 meningeal single cell transcriptome data set (DeSisto et al., 2020) and ran a sub-clustering analysis on cells expressing E-cadherin (*Cdh1*), which is only expressed by arachnoid barrier cells in the meninges. The analysis identified three subclusters at different putative stages of arachnoid barrier development that we called “Mature”, “PreAB-1” and “PreAB-2” (Figure 1A and 1B). Cells in the Mature AB subcluster had the highest expression of E-cadherin (*Cdh1)*, Claudin 11 (*Cld11*) and Klf5 (*Klf5)* (Figure 1B). Pseudo-time analysis using Slingshot (Street et al., 2018) identified a distinct hierarchical lineage relationship among the three arachnoid barrier subclusters (Figure 1A) and an Ingenuity Pathway Analysis identified Wnt-β-catenin signaling as significantly down regulated in the cells of the mature AB sub-cluster compared to both PreAB clusters (log10^−3^ adjusted P-value) (Figure 1C). This data sets up the prediction that the differentiation of arachnoid barrier cells from unspecified meningeal mesenchymal cells requires downregulation of Wnt-β-catenin signaling.

**Figure 1.**
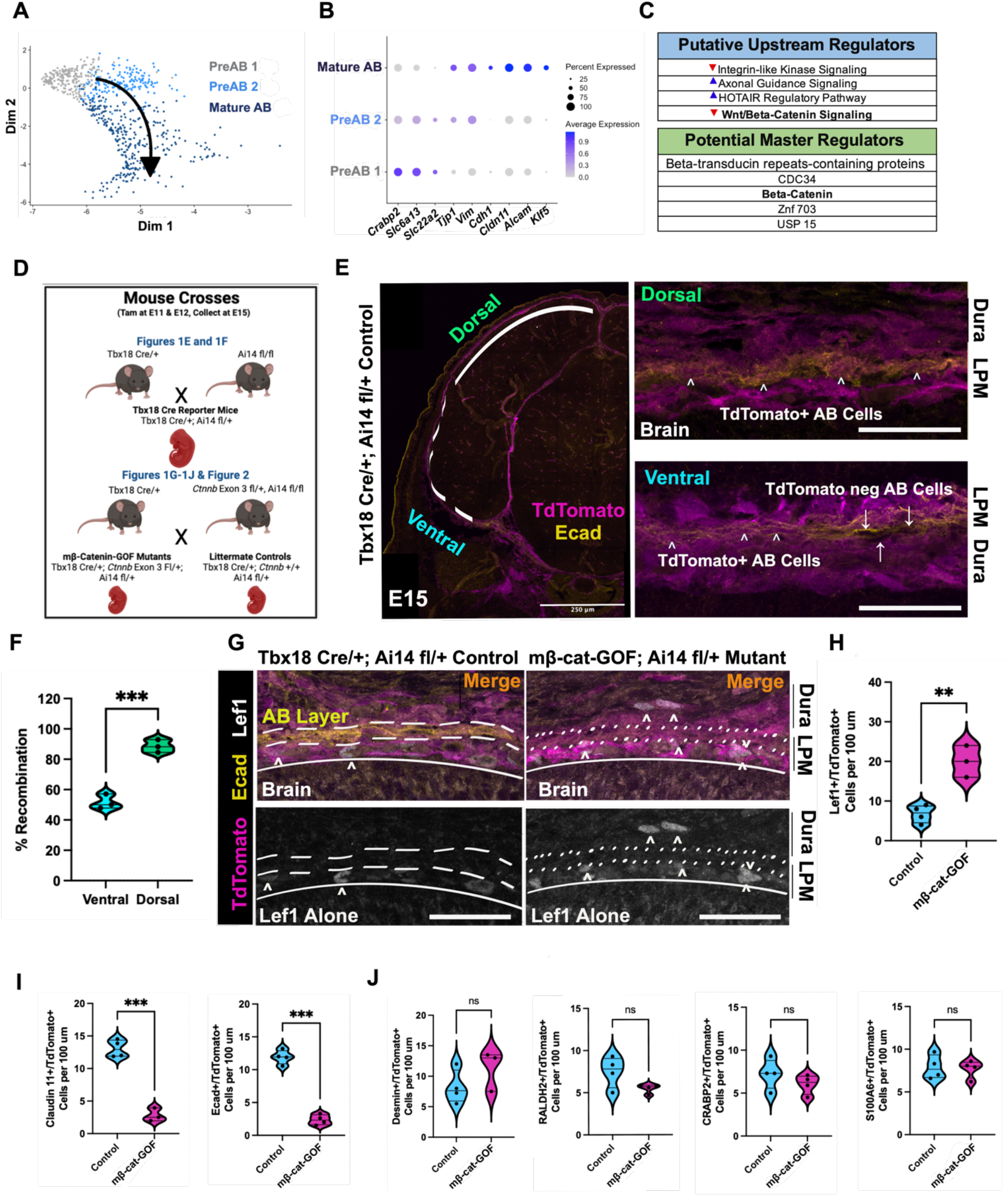
Downregulation of Wnt-B-catenin signaling is required for proper specification of arachnoid barrier cells. (A) UMAP plot of reclustered E14 meningeal cells that came from all clusters with *Cdh1+* cells from original analysis with arrow depicting the pseudotime lineage. (B) Dotplot of genes showing progression through pseudotime lineage with PreAB1 expressing arachnoid non-barrier cell marker *Crabp2* and two Slc transporters, whereas PreAB2 begins to upregulate *Tjp1* and *Vim* showing a transition towards arachnoid barrier cell identity and ‘Mature’ AB cells express *Tjp1, Vim, Cdh1, Cldn11, Alcam*, and *Klf5*. (C) Ingenuity Pathway Analysis results comparing the differentially expressed genes of PreAB1 and PreAB2 vs. Mature arachnoid barrier cells. (D) Schematic of mouse crosses for Figures 1E and 1 F (top) and Figures 1G-J and Figure 2 (bottom). All mice received tamoxifen at E11 and E12. (E) Low magnification image of E15 Tbx18-CreERT; Ai14-fl/+ labeled with E-cadherin with Dorsal (solid line and green labeling on left, top row on right) and Ventral (dashed line and teal labeling on left, bottom row on right) with carrots pointing to recombined TdTomato+ and E-cadherin+ arachnoid barriercells. Arrows indicate TdTomato-and E-cadherin+ arachnoid barrier cells that are notrecombined. (F) Quantification of recombination percentages in ventral and dorsal areas. (G)Representative images of Lef1, E-cadherin (representing the arachnoid barrier or AB layer), and TdTomato expression in the dorsal forebrain meninges of *Tbx18* Cre/+; Ai14-fl/+ Control vs. mβ-cat-GOF. Carets indicate Lef1+/tdTomato+ cells. Dotted line in mβ-cat-GOF indicate region of tdTomato+ leptomeninges (LPM) lacking E-cadherin+ arachnoid barrier layer. (H) Quantification of Lef1+/TdTomato+ cells per 100 μm. (I) Quantification of TdTomato+ arachnoid barrier cells, Claudin 11+ left and E-cadherin+ on right, per 100 μm. (J) Quantification of TdTomato+/Desmin+ (pericytes), TdTomato+/Raldh2+ (arachnoid fibroblasts), TdTomato+/Crabp2+ (arachnoid and dura fibroblasts), and TdTomato+/S100A6+ (pial fibroblasts). Statistics: Students T-Tests, *** = p < 0.01, *** = p < 0.001. Scale bars =250 μm in (E low magnification) and 50 μm in (E high magnification and G).

To test this prediction, we analyzed arachnoid barrier cell specification in mice where repression of Wnt signaling is prevented. We generated these mice to express three alleles: 1) the *Tbx18-CreERT2* allele, which drives Cre expression in all meningeal mesenchymal cells and mural cells, 2) a floxed β-catenin gain of function allele (*Ctnnb1-*ex3-flox) which will delete exon 3 on β-catenin, thereby preventing β-catenin to be phosphorylated and degraded upon Cre activation(Harada et al., 1999), and 3) a floxed tomato reporter allele (Ai14) (Figure 1D). To show that the *Tbx18-CreERT2* allele can drive recombination in arachnoid barrier cells during late embryogenesis, we injected *Tbx18-CreERT2; Ai14-Flox* expressing mice with tamoxifen at E11 and E12, a time point prior to arachnoid barrier cell specification (DeSisto et al., 2020) and collected E15 whole head. At E15, E-cadherin+ arachnoid barrier cells are present in a continuous layer that covers the entire ventral to dorsal surfaces of the forebrain (Fig. 1E). Cre induced tdTomato expression was strong in E-cadherin+ arachnoid barrier cells, showing ∼90% expression rate in dorsal arachnoid barrier cells and lower, ∼50% recombination rate in ventral arachnoid barrier cells (Figure 1F). Thus, we focused our analyses on the dorsal forebrain meninges with high recombination rates.

To constitutively elevate Wnt-β-catenin signaling in meningeal cells prior to arachnoid barrier cell specification, we exposed females with *Tbx18-CreERT2; Ctnnb1-*ex3-flox/+; *Ai14 flox/+* embryos (meningeal β-catenin-GOF or mβ-cat-GOF) to tamoxifen at E11 and E12. At E15, mβ-cat-GOF had increased expression of direct Wnt-β-catenin target Lef1 in tdTomato+ cells as compared to littermate controls (*Tbx18-CreERT2; Ai14-fl/+*) (Figure 1G and 1H), confirming elevated Wnt signaling in recombined cells. Claudin11+ and E-cadherin+ tdTomato+ cells were nearly absent from the mβ-cat-GOF E15 dorsal forebrain meninges (Figure 1G and 1I), demonstrating substantial failure of arachnoid barrier cell development in these mutants. Elevated Wnt-β-catenin signaling did not alter the density of other meningeal cell types recombined by *Tbx18-CreERT2*, including desmin+ mural cells, Raldh2+ arachnoid fibroblasts, CRABP2+ pan-arachnoid and dura fibroblasts, or S100A6+ pial fibroblasts cells in the mβ-cat-GOF E15 dorsal forebrain meninges (Figure 1J). Heads and brains of E17 mβ-cat-GOF were grossly normal (Supp. Figure 1), with no obvious neocortical defects. This is important to note because embryos with more substantial meningeal developmental defects have accompanying brain abnormalities(Siegenthaler et al., 2009; Zarbalis et al., 2007). Collectively this shows that arachnoid barrier cell specification requires a repression of Wnt-β-catenin signaling.

To test the functional relevance of the fetal arachnoid barrier, we investigated the permeability of the wildtype and mβ-cat-GOF meninges to a small molecular weight fluorescent tracer (Biocytin-TMR, 893 Daltons) and the leading meningeal bacterial pathogen, GBS (Figure 2A). Ten minutes after an injection of biocytin-TMR, we removed the meninges, dissected a piece of superficial cortex to quantify tracer content by plate reader and discovered that there was a significant increase in brain biocytin-TMR content in E17 mβ-cat-GOF mice as compared to littermate controls (Figure 2C). We imaged whole head sections and found that mβ-cat-GOF mutants had increased biocytin-TMR only in regions near the cortical surface of the dorsal forebrain, but not in the deeper subventricular zone away from the brain surface (Figure 2B, 2D, 2E). This supports that the BBB is intact in mβ-cat-GOF mutants and biocytin-TMR is ‘leaking’ into the brain due to an absent arachnoid barrier layer in this area. We further visualized and analyzed the biocytin-TMR signal in the leptomeninges (pia and arachnoid) and observed that while both control and mβ-cat-GOF mutant mice had a high tracer fluorescent intensity in the meninges at 7-8 microns away from the brain surface (where controls have Claudin11+ arachnoid cells, but mβ-cat-GOF lack arachnoid barrier cells), there was a precipitous drop off in tracer signal in the pia and arachnoid in the controls. In contrast, biocytin-TMR content remained high in the mβ-cat-GOF mutants throughout the entire leptomeninges and down to the brain surface (Figure 2B and 2F). This is strong evidence that the arachnoid barrier is functional by E17 and the lack of an arachnoid barrier in mutants permits access of peripheral molecules into the fetal leptomeninges and brain.

**Figure 2.**
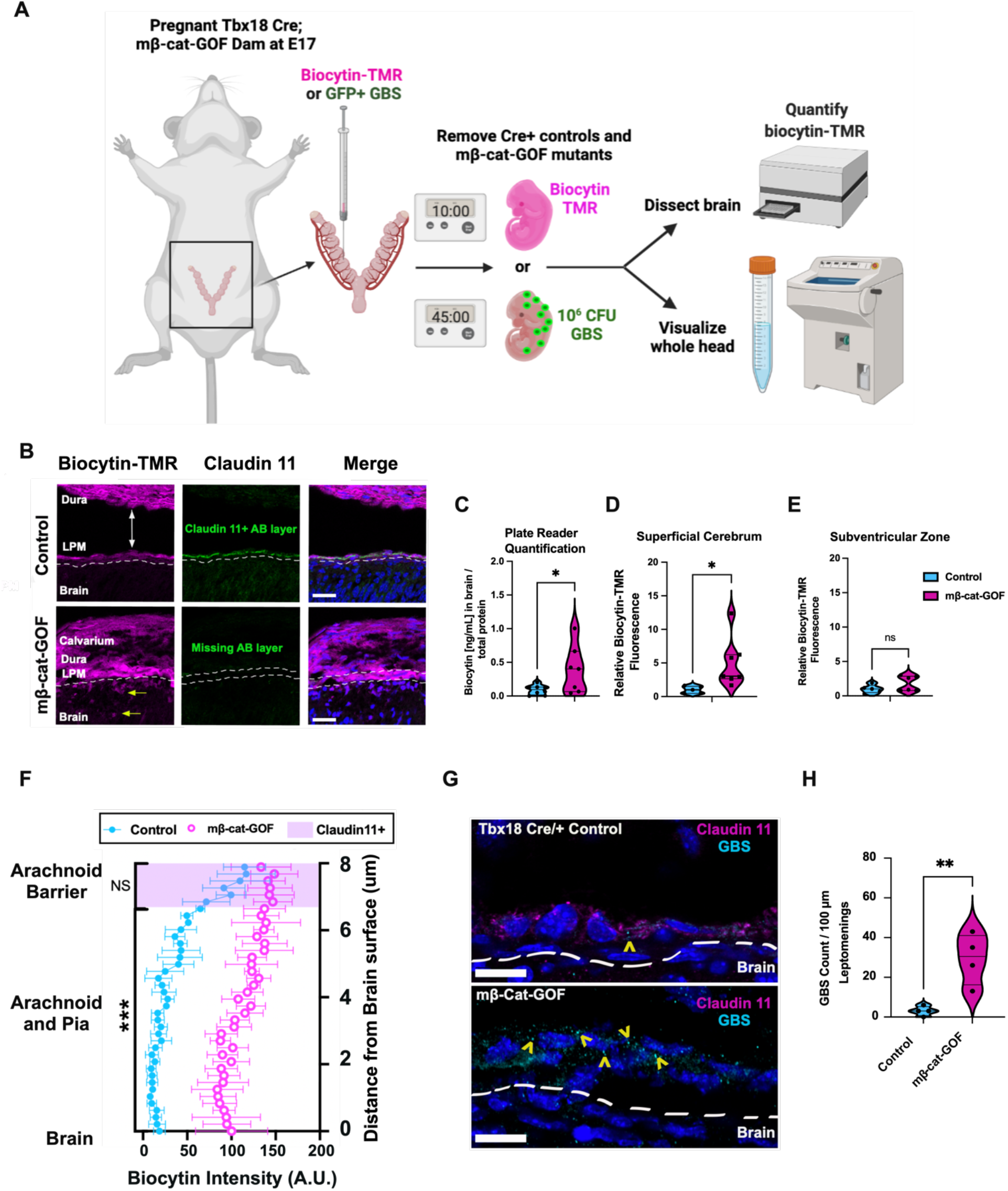
The arachnoid barrier is critical to prevent unwanted molecules and pathogens from entering the CNS. (A) Schematic of experimental design showing E17 embryos from Tbx18Cre; mβ-cat-GOF dams receiving either 20uL of Biocytin-TMR to circulate for 10 minutes or 20 µL of 10^6^ C.F.U. of GFP+ GBS to circulate for 45 minutes. After injections, biocytin animals were sacrificed and brain were dissected, homogenized, and read on a plate reader or whole heads were fixed, sectioned, and visualized via immunohistochemistry and microscopy. GBS infected animals, the whole heads were dissected for sectioning and visualization by microscopy. (B) Representative images of N=5 control and N=7 mβ-cat-GOF mutants. Yellow single-headed arrows indicate cells with tracer in them inside of the brain. White double-headed arrow indicates where the leptomeninges and dura (attached to the calvarium) separate in the control due to a tissue processing artifact in control samples. (C) Quantitation of biocytin-TMR in brain, normalized to total protein, from the plate reader assay. (D) Quantification of biocytin-TMR fluorescence intensity from images at the superficial cerebrum (50 microns from pial surface). (E) Quantification of biocytin-TMR fluorescence intensity from images in cerebral subventricular zone (50 microns from lateral ventricle surface). (F) Line analysis of leptomeninges (Claudin 11+ is shaded in pink to indicate where Claudin11+ arachnoid barrier cells are in the control animals) and descending 8 microns from arachnoid barrier cells, through the arachnoid and pia. (G) Representative images of N=4 mβ-cat-GOF and N=4 control animals infected with GBS-GFP. Yellow carrots point to GFP+ GBS puncta in the leptomeninges and brain. (H) Quantification of images to determine GBS burden per 100 μm of dorsal meninges. Statistics: Students T-Tests for 2C, 2D, 2E, 2F, and 2G and 2H, with average of all arachnoid and pia intensity values collapsed into one value for 2F, * = p < 0.05, ** = p < 0.01, *** = p < 0.001. Scale bars represent 50 μm in B and 20 μm in G.

To further test the functional relevance of the arachnoid barrier, we infected mβ-cat-GOF mutant fetuses and littermate controls across two litters with GBS, a physiologically relevant pathogen that drives bacterial meningitis in neonates, to determine if an impaired arachnoid barrier altered pathology. We injected E17 fetuses in the liver with 10^6^ Colony Forming Units (C.F.U.) of GFP-labeled GBS (∼1 µm in diameter) and collected tissue 45 min later (Figure 2A). While GFP-GBS+ in leptomeninges of the littermate control animals were sparse, consistent with an intact, functional arachnoid barrier, we observed a significantly more severe GBS burden in the leptomeninges of the mβ-cat-GOF mutants, with no major differences in GBS burden in skin (Supplemental Figure 2). This indicates that the lack of an arachnoid barrier in mβ-cat-GOF mutants renders these animals increasingly susceptible to GBS infiltration into the CNS (Figure 2G and 2H).

To further detail the maturation of the arachnoid barrier we sought to determine the age when mature E-Cadherin/Claudin 11 double positive arachnoid barrier cell can be detected over the entirety of the brain and determine the earliest age when the arachnoid barrier can functionally restrict the passage a small molecule tracer. At E12, E-Cadherin/Claudin 11 double positive arachnoid barrier cells are not detectable in the meninges (Fig 3A 3B). Note: a few cells are expressing cytosolic Claudin 11 at E12 in/near the brain and spinal cord, but these cells do not co-express E-cadherin, indicative of another cell type (Figure 3A and 3B). At E13 we observed consistent E-cadherin and Claudin11 double positive cells forming a layer in the ventral hindbrain with a patchy layer of a double-positive cells in the ventral forebrain, but very few double positive cells in the dorsal forebrain (Figure 3A-B). By E14, the number of double positive cells in the dorsal forebrain had increased, which forms a continuous layer in all regions by E15 (Figure 1E).

**Figure 3.**
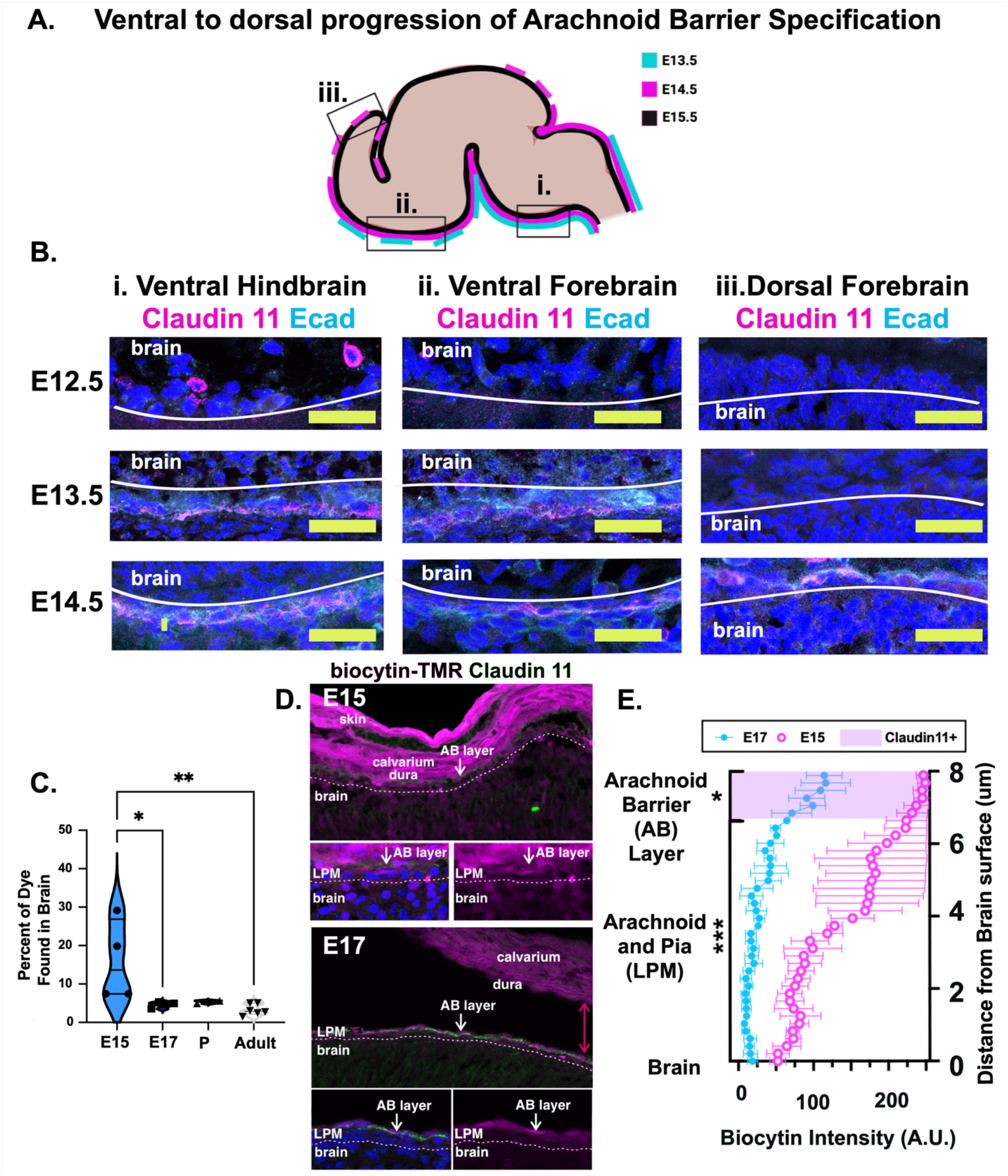
Arachnoid barrier cells have a ventral to dorsal wave of specification from E13-E15, which precedes barrier formation by E17. (A) Schematic of embryonic murine brain with the timing that Claudin 11 and E-cadherin are patchy (dashed line) or contiguous (solid line) around the brain with boxes at i. Ventral Hindbrain, ii. Ventral Forebrain, and iii. Dorsal Forebrain regions where images in B were taken. (B) Representative images of sections with immunolabeling for Claudin11 (magenta) and E-cadherin (cyan) from various locations indicated in A, showing progressing appearance of arachnoid barrier cells from E12-E15. (C) Quantification of biocytin-TMR assay in brain across ages when N=4 animals were injected with 0.25% Biocytin-TMR. (D) Representative images of whole heads at the dorsal forebrain, E15 and E17 biocytin-TMR injected embryos immunolabeled with Claudin 11 to show location of arachnoid barrier (AB) layer. (E) Quantification of tracer fluorescence within the leptomeninges of E15 vs. E17 embryos from Claudin11+ arachnoid barrier cells 8 microns above the brain surface and down through the lower arachnoid and pia. Two-way Students T-Tests for 3C and 3D, with average of all arachnoid barrier or arachnoid and pia intensity values collapsed into one value for 3D, * = p < 0.05, ** = p < 0.01, *** = p < 0.001. Scale Bars are 50 μm in B.

We next tested if the presence of a continuous layer at E15 resulted in a functional barrier by performing biocytin-TMR assays at E15 as described in Figure 2. Plate reader analysis showed that the superficial cortex of E15 brains had significantly more fluorescent signal than the superficial cortexes of brains from E17, postnatal, and adult animals (Figure 3C). Analysis of tissue sections confirmed these findings showing that the amount of dye below the arachnoid barrier, in the leptomeninges and superficial brain, was significantly higher in the E15 embryos compared to E17 embryos (Figure 3D, E). Collectively, these analyses demonstrate that the arachnoid barrier cellular layer is grossly continuous by E15, but the functional ability to restrict the passage of small molecular weight tracers into the leptomeninges does not occur until later.

We next sought to detail the maturation of tight and adherens junctions between arachnoid barrier cells, since tissue barrier integrity critically dependent on mature junctions. Consistent with electron microscopy and freeze fracture studies (Balin et al., 1986; Nabeshima et al., 1975), adult cerebral leptomeninges whole mounts show arachnoid barrier cells are connected by tight and adherens junctions marked by co-localization of E-cadherin, Claudin 11 and ZO-1 proteins (Figure 4A). To identify when junctions appear between arachnoid barrier cells and how junctional localization changes during growth of a continuous arachnoid barrier layer from E15 to adult, we used leptomeninges wholemounts (Figure 4B). These are ideal to analyze arachnoid barrier layer junctional protein localization (junctional vs non-junctional) and junctional density at E15, E17, P4, P7 and adults (Bonney et al., 2019; Hannah E. Jones et al., 2022) (Figure 4C and 4D). At E15, when the arachnoid barrier layer has not yet developed the functional ability to restrict small molecule tracers, Claudin 11 protein expression is diffuse and mostly non-junctional (Figure 1B) with fewer Claudin 11+ junctions compared to E-cadherin junctions (Figure 4G). E15 E-cadherin junctions were more abundant, but had lower junctional fluorescence intensity as compared to later time points (Figure 4F – 4G). By E17, the number of Claudin 11+ and E-cadherin+ junctions had increased and reached equal numbers (Figure 4B and 4G). We also observed from pre-to postnatal time points, there was a decrease in non-junctional expression of Claudin 11 (Figure 4B, E) and to a lesser extent E-cadherin (Figure 4B, F). Collectively this analysis shows that acquisition of a functional barrier coincides with elevated junctional localization of Claudin 11 and increased E-cadherin.

**Figure 4.**
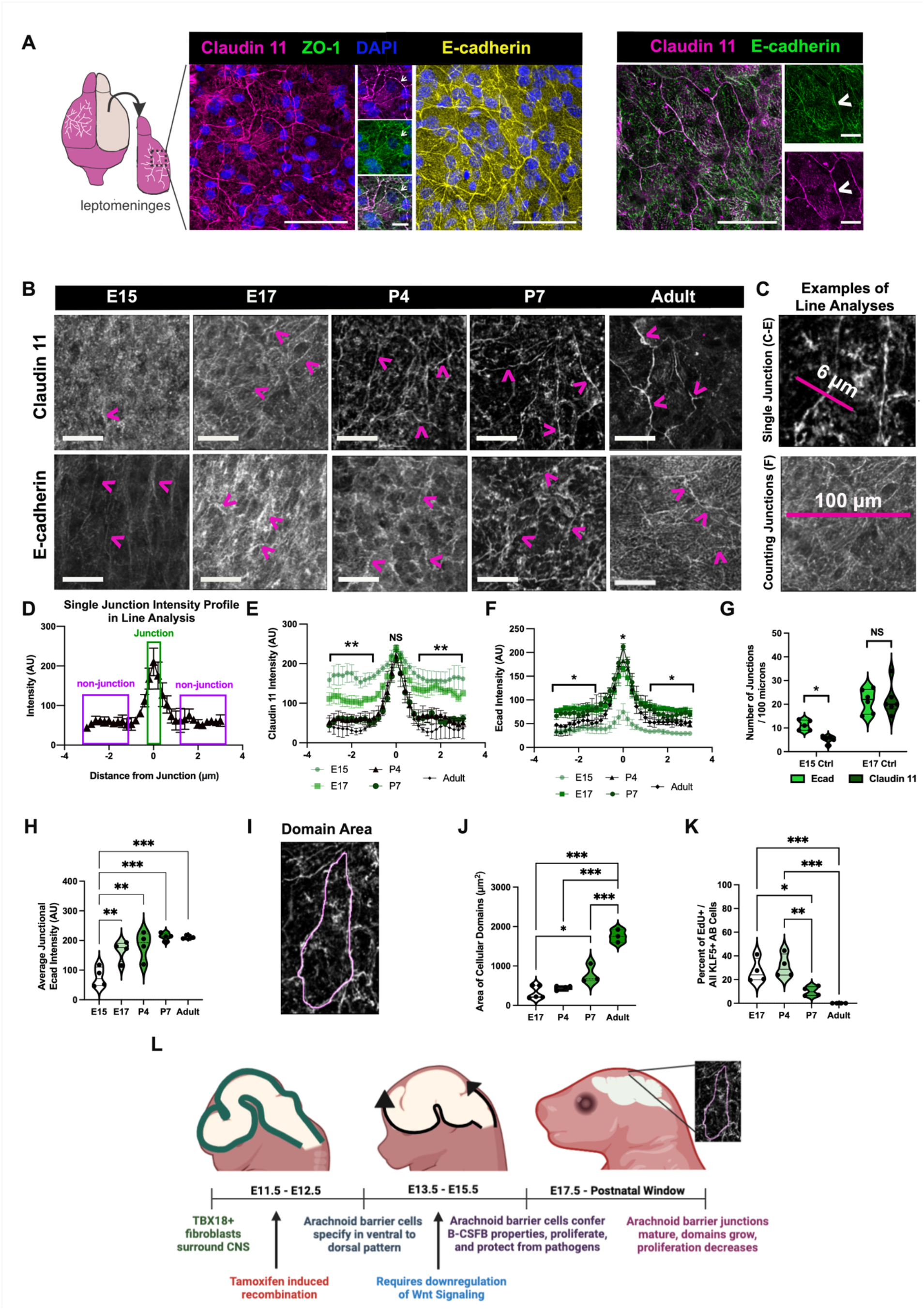
Embryonic emergence of arachnoid barrier tight and adherens junctions that mature postnatally. (A) Diagram and representative images of adult cerebral leptomeninges wholemounts to visualize tight (Claudin 11, ZO-1) and adherens (E-cadherin) junctions proteins that connect arachnoid barrier cells. Co-localization of Claudin l1/ZO-1 and Claudin 11/E-cadherin is observed at major junctions between adjacent cells. (B) Representative images of cerebral leptomeninges whole mounts harvested from wildtype mice at E15, E17, P4, P7, and adult timepoints immuolabled with Claudin 11 (top row) and E-cadherin (bottom row). Arrows are pointing at junctions that were quantified. (C) Examples of quantification approach for a single tight junction (top) for 4E and 4F and example for quantification approach for counting the number of junctions per 100 μm for 4G (bottom). (D) Example data from single junction intensity profile analysis where non-junctional intensity was quantified as 1-3 μm away from the junction on either side and junctional intensity was quantified as the central peak in the analysis. (D) Quantification of Claudin11+ single junction line intensity from E15 to adult. (E) Quantification of E-cadherin+ single junction line intensity from E15 to adult. (G) Number of junctions found in E15 and E17 leptomeningeal whole mounts per 100 μm, as defined by a peak that is >50 florescence AU higher than the fluorescence values 1-2 μm away for E-cadherin and Claudin 11. (H) Average junction intensity of Ecad from E15 to Adult. (I) Example of how domain area was quantified. (J) Increasing area of cellular domains in Claudin11+ arachnoid barrier cells from E17-Adult. (K) Quantification of the percent of proliferating cells after a 2h EdU pulse as defined by EdU+/Klf5+ cells divided by all Klf5+ arachnoid barrier cells from E17-Adult. (K) Schematic showing developmental timeline of arachnoid barrier specification and maturation. Statistics: Students T-Tests for 4E-G (all non-junctional values collapsed into 1 value for 4E and 4F) and Two-way ANOVA for 4H, 4J, 4K. * = p < 0.05, ** = p < 0.01, *** = p < 0.001. Scale bar = 50 μm in A, B and 20 μm in insets in A.

Arachnoid barrier cells are connected by junctions on multiple sides, creating domains. We observed that arachnoid barrier domain area increased from E17 to P7 and adult time points (Figure 4I, J). Arachnoid barrier cell proliferation measured by EdU pulse-labeling reduced over these age ranges (Figure 4K), indicative of a concomitant decrease in proliferation as domain size increases during brain growth.

Taken together, a model of arachnoid barrier layer development emerges from our data. First, the arachnoid barrier cell layer arises from meningeal mesenchymal precursors in a temporally and spatially restricted manner from E13 to E15 through a mechanism involving downregulation of Wnt-β-catenin. In the cerebral leptomeninges, E-cadherin forms adherens junctions between arachnoid barrier cells by E15, followed by junctional Claudin 11 and a functional barrier by E17. To accommodate brain surface area growth from prenatal to adult stages, the arachnoid barrier cells proliferate in pre-and early postnatal stages, but later arachnoid barrier domain areas increase to expand the arachnoid barrier layer while maintaining a functional barrier (Figure 4L). These mechanisms work in concert to protect the CNS from peripheral infiltration of molecues and pathogens as early as E17 and throughout the lifespan.

## Discussion

The arachnoid barrier is a fundamental component of the B-CSFB. A small number of structural and tracer leakage studies support a functional role as a CNS barrier (Balin et al., 1986; Nabeshima et al., 1975; Rascher and Wolburg, 1997; Rodriguez-Peralta, 1957). However, there have been no reports to describe how the arachnoid barrier forms, when it becomes functional, or the clinical implications when arachnoid barrier integrity is absent or impaired. This study provides the first insights into the cellular and functional phenotypes during arachnoid barrier formation and describes arachnoid barrier cells from their first emergence at E13, to the materialization of tight junctions and barrier properties by E17 and through the postnatal maturation process as the arachnoid barrier layer expands around the growing brain. This process is similar to the prenatal emergence and postnatal maturation of the BBB, with key differences. The arachnoid barrier forms just after the BBB which is restrictive to tracers by E13-E14, depending on brain region (Ben-Zvi et al., 2014). Arachnoid barrier cells are specified *de novo* from meningeal mesenchymal cells, dependent on downregulation Wnt-β-catenin signaling. This is in contrast to BBB vasculature which grows into the CNS via angiogenesis of existing vasculature surrounding the CNS and acquires barrier properties via upregulation of Wnt-β-catenin signaling, induced by locally produced Wnt ligands (Daneman et al., 2009; Liebner et al., 2008; Stenman et al., 2008; Zhou et al., 2014).

These studies also give a glimpse into the critical contributions of the arachnoid barrier in protecting the CNS, particularly against susceptibility to GBS invasion. These findings parallel previous work that indicates the postnatal maturation of the choroid plexus junctional integrity is critical to protect against neonatal infections (Travier et al., 2021). Our work modeling the timing of arachnoid barrier development provides a framework to better understand how and why neonatal babies, potentially with an arachnoid barrier that is still in a dynamic cellular state of maturation due to ongoing brain growth, may be uniquely vulnerable to CNS access of certain pathogens. Importantly, our model of defective arachnoid barrier development, whole mounts to visualize arachnoid barrier layer junctional organization, and functional assays may aid in studies to address outstanding questions about the arachnoid barrier. These include the basic biology of the arachnoid barrier. How does Wnt-β-catenin signaling suppression support arachnoid barrier cell specification and how are E-cadherin and Claudin 11 localized to cell-cell junctions in newly specified arachnoid barrier cells to form a functional structure? Disease relevant questions include how the arachnoid barrier responds to acute physical injury vs. infectious agents vs. chronic disease, and can the arachnoid barrier reactivate developmental programs to allow for regeneration.

## Methods and Materials

### Mice

Mice used for experiments here were housed in specific-pathogen-free facilities approved by the AALAC and were handled in accordance with protocols approved by the University of Colorado Anschutz Medical Campus IACUC committee. C57/Bl6 (Jackson Laboratory). The following mouse strains were used in this study: 1) Tbx18^tm3.1(cre/ERT2)Sev/J^ (Jackson Laboratory Stock No: 031520; RRID: IMSR_JAX:031520), 2) Ai14fl/fl (Gt(ROSA)26Sor^tm14(CAG-tdTomato)Hze,^ Jackson Laboratory Stock No: 007914 RRID: IMSR_JAX:007914) and β-cat-GOF *Ctnnb1*^lox(ex3)^ and 3) C57Blk/6J (Jackson: 000664) wildtype animals were bred to generate embryonic or postnatal litters (E15, E17, P4, P7) or collected as adults (12 weeks) for wholemounts analysis. For timed pregnancy, males and females were placed together on the afternoon and presence of a plugged was checked AM daily; the day the plug was detected was counted as embryonic day 0.5 To activate Cre-mediated recombinase activity, pregnant dams were injected intraperitoneally with 100 μl of tamoxifen (Sigma, Cat#: T5648) dissolved in corn oil (Sigma cat#: C8267; 20 mg/ml) at embryonic day 11 and 12. To measure arachnoid barrier tracer leakage, embryonic mice were injected in liver with 20 μl of 0.25% Biocytin-TMR (Thermo Fisher) for 10 min prior to isolation of tissue for immunohistochemistry or plate reader analysis at their specific timepoint. Postnatal and adult mice were injected intraperitoneally 30 min prior to isolation of tissue, were postnatal mice were injected with 25 μl of 0.25% Biocytin-TMR and adult mice were injected with 100 μl of 0.25% Biocytin-TMR, with a protocol adapted from the Gu Laboratory for BBB analysis(Ben-Zvi et al., 2014). For GBS inoculation, mice were injected in the liver with ∼1.2 × 10^6^ CFU and sacrificed 45 min later for isolation of tissues. For EdU detection, we administered EdU (2.5 mg/mL) via intraperitoneal injection prior to anesthetization (embryonic: 150 µL into pregnant dam, P0-P21: 50 µL, adult: 100 µL) and allowed to circulate for 2 h prior to anesthetization.

### Bacterial Culture

*Streptococcus agalactiae*, (Group B *Streptococcus*, GBS) strain COH1+pDESTerm-GFP(Mu et al., 2014), has been described previously described(Mu et al., 2014) grown statically overnight at 37°C in Todd-Hewitt Broth supplemented with 5 µg/mL erythromycin. Bacteria were sub-cultured and grown to mid-exponential phase before resuspension in PBS.

### Bioinformatic Analysis of Single Cell Data

R Studio with R version 3.6.3 and Seurat V.4 were used for further analysis of our previously published dataset (DeSisto et al., *Developmental Cell*, 2020)(DeSisto et al., In Press). All clusters that contained arachnoid barrier cells, as defined by *Cdh1* at >1 expression, were subsetted out and subclustered using FindNeighbors(reduction=‘pca’, dims=1:30) and FindClusters(resolution=0.5). Slingshot pseudotime lineage analysis was applied to the new subclusters to determine the lineage pathway from immature to mature arachnoid barrier cells(Street et al., 2018). Subsequently, Ingenuity Pathway Analysis (Qiagen) was applied to lists of differentially expressed genes between each cluster to determine putative upstream and master regulators of the changes seen across the trajectory.

### Immunohistochemistry of Whole Head Sections

Embryos were collected and whole heads fixed for 48h with 4% paraformaldehyde followed by 20% sucrose and frozen in OCT compound (Tissue-Tek). Tissue was cryosectioned in 12μm increments and tissue-mounted slides were subjected to antigen retrieval by immersing the slides in solution of 0.01M citric acid, 0.05% Tween, pH 6, and heating in a pressure cooker (Cuisinart Model CPC-600) for 6 min. No antigen retrieval was performed when staining for Claudin11, Desmin, or E-cadherin. The tissue was permeabilized by incubating for 10 min at room temperature in PBS with 0.1% Triton-X (Sigma), blocked in 2% lamb serum/0.05% Triton-X solution for 40 min at room temperature and primary and secondary solution was 0.05% Triton-X in PBS. Incubation in the following primary antibodies was conducted overnight at 4°C in appropriate solution: rabbit anti-S100A6 (1:100; Novus NBP2-44492), mouse anti-CRABP2 (1:100 Millipore; MAB5488), rabbit anti-RALDH2 (1:100, Sigma-Aldrich HPA010022), chicken anti-GFP (1:500, Invitrogen A10262), mouse anti-E-cadherin (1:100, BD Transduction Laboratories 610181), rabbit anti-Desmin (1:100, Cell Signaling (5332S), rabbit anti-Lef1 (1:100, Cell Signaling 2230S, and rabbit anti-Claudin 11 (1:100, Thermofisher, PIPA568608), and chicken anti-RFP (1:200; Rockland). Following incubation with primary antibodies, tissue sections were incubated for 60 min with appropriate Alexafluor-conjugated secondary antibodies (Invitrogen) and DAPI (1:1000; Invitrogen).

### Whole Mount Preparations and Immunohistochemistry

All whole mount dissection and immunostaining and EdU detection in whole mount are detailed at length in our protocol paper Jones et al., 2022 *Neurophotonics*(Hannah E. Jones et al., 2022).

### Microscopy Imaging and Image Quantification

Confocal images were obtained using a Zeiss 780 Laser Scanning Microscope with associated Zeiss Zen Black software. Images were processed in FIJI Image J and Graphic software. For cell counts in Figure 1, images were subjected to thresholding and the number of positive cells were counted per 100 μm of leptomeninges was counted (or 100 μm of E-cadherin+ cells, if listed). For Figure 2D, a 15 μm line was drawn in FIJI within 20 μm of the cortical surface or where nuclei patterns in DAPI indicated the subventricular zone of the mouse brain and the average intensity across that line was recorded for each sample. For Figure 1E, 5 control E17 animals with Claudin 11 staining were used to determine the average thickness of the arachnoid barrier (∼1.7 μm) and the distance from the outer surface of the arachnoid barrier to the pial/brain surface (∼8 microns). Subsequently, a line from the pial/brain surface was extended up for 8 microns for both control and mutant animals and the biocytin intensity was plotted spatially. For Figure 2G, images were separated by color with the green (488 nm) channel subjected to thresholding, and a box of 8 μm (height of the arachnoid barrier) by 100 μm was drawn to count all GFP+ GBS (> 0.5 μm in diameter). For Figures 4D and 4E, a 6 μm line was drawn across the junctions (as depicted in 4B) and an intensity profile was plotted with the outer 2.5 μm indicating cytosolic portions of the cell and the middle peak indicating the junction. For Figure 4H, the perimeter of cell domains was traced using the freehand tool in FIJI and the area within was measured for each sample. E15 were excluded from this analysis because it was not possible to determine a unique cellular domain at this age. For 4I, KLF5 was used to mark arachnoid barrier nuclei and EdU+/KLF5+ nuclei were quantified as proliferating arachnoid barrier cells. Data is plotted as a percentage, quantified by the division of EdU and KLF5 double positive nuclei by all KLF5 positive nuclei * 100.

## Statistical analysis

Statistical analyses were performed as described in the text using GraphPad Prism, where Two-Way ANOVAs and multiple comparisons with Tukey adjustments were performed in instances with two or more groups and Students’ T-tests were performed to compare groups of two, after normality assumption was achieved.

## Acknowledgements

This work was supported by RO1 NS098273 to J.A.S. from NIH/NINDS, F32 NS122999 for J.D. from the NIH/NINDS, and K.S.D laboratory R01NS116716 from the NIH/NINDS.

## Notes

### Competing Interest Statement

The authors have declared no competing interest.

## REFERENCES

Alves de Lima, K., Rustenhoven, J., and Kipnis, J. (2020). Meningeal Immunity and Its Function in Maintenance of the Central Nervous System in Health and Disease. Annu. Rev. Immunol. 38, 597–620. https://doi.org/10.1146/annurev-immunol-102319-103410.

Balin, B.J., Broadwell, R.D., Salcman, M., and El-Kalliny, M. (1986). Avenues for entry of peripherally administered protein to the central nervous system in mouse, rat, and squirrel monkey. J. Comp. Neurol. 251, 260–280. https://doi.org/10.1002/cne.902510209.

Ben-Zvi, A., Lacoste, B., Kur, E., Andreone, B.J., Mayshar, Y., Yan, H., and Gu, C. (2014). MSFD2A is critical for the formation and function of the blood brain barrier. Nature 509, 507–511. https://doi.org/10.1038/nature13324.

Bonney, S., Seitz, S., Ryan, C.A., Jones, K.L., Clarke, P., Tyler, K.L., and Siegenthaler, J.A. (2019). Gamma Interferon Alters Junctional Integrity via Rho Kinase, Resulting in Blood-Brain Barrier Leakage in Experimental Viral Encephalitis. MBio 10, e01675–19. https://doi.org/10.1128/mBio.01675-19.

Brkic, M., Balusu, S., Van Wonterghem, E., Gorle, N., Benilova, I., Kremer, A., Van Hove, I., Moons, L., De Strooper, B., Kanazir, S., et al. (2015). Amyloid Oligomers Disrupt Blood-CSF Barrier Integrity by Activating Matrix Metalloproteinases. J. Neurosci. 35, 12766–12778. https://doi.org/10.1523/JNEUROSCI.0006-15.2015.

Brøchner, C.B., Holst, C.B., and Møllgård, K. (2015). Outer brain barriers in rat and human development. Front. Neurosci. 9, 75–75. https://doi.org/10.3389/fnins.2015.00075.

Daneman, R., Agalliu, D., Zhou, L., Kuhnert, F., Kuo, C.J., and Barres, B.A. (2009). Wnt/beta-catenin signaling is required for CNS, but not non-CNS, angiogenesis. Proc. Natl. Acad. Sci. U. S. A. 106, 641–646. https://doi.org/10.1073/pnas.0805165106.

Derk, J., Jones, H.E., Como, C., Pawlikowski, B., and Siegenthaler, J. (2021). Living on the edge of the CNS: meninges cell diversity in health and disease. Front. Cell. Neurosci. Accepted-In Press..

DeSisto, J., O’Rourke, R., Jones, H.E., Pawlikowski, B., Malek, A.D., Bonney, S., Guimiot, F., Jones, K.L., and Siegenthaler, J.A. (2020). Single-Cell Transcriptomic Analyses of the Developing Meninges Reveal Meningeal Fibroblast Diversity and Function. Dev. Cell 54, 43–59.e4. https://doi.org/10.1016/j.devcel.2020.06.009.

DeSisto, J., O’Rourke, R., Bonney, S., Jones, H.E., Guimiot, F., Jones, K.L., and Siegenthaler, J.A. (In Press). A cellular atlas of the developing meninges reveals meningeal fibroblast diversity and function. Dev. Cell 648642. https://doi.org/10.1101/648642.

Hannah E. Jones, Kelsey A. Abrams, and Julie A. Siegenthaler (2022). Techniques for visualizing fibroblast-vessel interactions in the developing and adult CNS. Neurophotonics 9, 1–17. https://doi.org/10.1117/1.NPh.9.2.021911.

Harada, N., Tamai, Y., Ishikawa, T., Sauer, B., Takaku, K., Oshima, M., and Taketo, M.M. (1999). Intestinal polyposis in mice with a dominant stable mutation of the beta-catenin gene. EMBO J. 18, 5931–5942. https://doi.org/10.1093/emboj/18.21.5931.

Ichikawa, H., and Itoh, K. (2011). Blood–arachnoid barrier disruption in experimental rat meningitis detected using gadolinium-enhancement ratio imaging. Brain Res. 1390, 142–149. https://doi.org/10.1016/j.brainres.2011.03.035.

Ichikawa, H., Ishikawa, M., Fukunaga, M., Ishikawa, K., and Ishiyama, H. (2010). Quantitative evaluation of blood–cerebrospinal fluid barrier permeability in the rat with experimental meningitis using magnetic resonance imaging. Brain Res. 1321, 125–132. https://doi.org/10.1016/j.brainres.2010.01.050.

Iliff, J.J., Wang, M., Liao, Y., Plogg, B.A., Peng, W., Gundersen, G.A., Benveniste, H., Vates, G.E., Deane, R., Goldman, S.A., et al. (2012). A paravascular pathway facilitates CSF flow through the brain parenchyma and the clearance of interstitial solutes, including amyloid β. Sci. Transl. Med. 4, 147ra111. https://doi.org/10.1126/scitranslmed.3003748.

Liebner, S., Corada, M., Bangsow, T., Babbage, J., Taddei, A., Czupalla, C.J., Reis, M., Felici, A., Wolburg, H., Fruttiger, M., et al. (2008). Wnt/beta-catenin signaling controls development of the blood-brain barrier. J. Cell Biol. 183, 409–417. https://doi.org/10.1083/jcb.200806024.

Liebner, S., Dijkhuizen, R.M., Reiss, Y., Plate, K.H., Agalliu, D., and Constantin, G. (2018). Functional morphology of the blood-brain barrier in health and disease. Acta Neuropathol. (Berl.) 135, 311–336. https://doi.org/10.1007/s00401-018-1815-1.

Mu, R., Kim, B.J., Paco, C., Del Rosario, Y., Courtney, H.S., and Doran, K.S. (2014). Identification of a group B streptococcal fibronectin binding protein, SfbA, that contributes to invasion of brain endothelium and development of meningitis. Infect Immun 82, 2276–2286. https://doi.org/10.1128/iai.01559-13.

Nabeshima, S., Reese, T.S., Landis, D.M.D., and Brightman, M.W. (1975). Junctions in the meninges and marginal glia. J. Comp. Neurol. 164, 127–169. https://doi.org/10.1002/cne.901640202.

Polanco, J., Reyes-Vigil, F., Weisberg, S.D., Dhimitruka, I., and Brusés, J.L. (2021). Differential Spatiotemporal Expression of Type I and Type II Cadherins Associated With the Segmentation of the Central Nervous System and Formation of Brain Nuclei in the Developing Mouse. Front. Mol. Neurosci. 14, 25. https://doi.org/10.3389/fnmol.2021.633719.

Rascher, G., and Wolburg, H. (1997). The tight junctions of the leptomeningeal blood-cerebrospinal fluid barrier during development. J. Hirnforsch. 38, 525–540..

Rodriguez-Peralta, L.A. (1957). The role of the meningeal tissues in the hemato-encephalic barrier. J. Comp. Neurol. 107, 455–473. https://doi.org/10.1002/cne.901070308.

Roth, TheodoreL., Nayak, D., Atanasijevic, T., Koretsky, AlanP., Latour, LawrenceL., and McGavern, DorianB. (2013). Transcranial amelioration of inflammation and cell death after brain injury. Nature 505, 223–228. https://doi.org/10.1038/nature12808.

Siegenthaler, J.A., Ashique, A.M., Zarbalis, K., Patterson, K.P., Hecht, J.H., Kane, M.A., Folias, A.E., Choe, Y., May, S.R., Kume, T., et al. (2009). Retinoic Acid from the Meninges Regulates Cortical Neuron Generation. Cell 139, 597–609. https://doi.org/10.1016/j.cell.2009.10.004.

Stenman, J.M., Rajagopal, J., Carroll, T.J., Ishibashi, M., McMahon, J., and McMahon, A.P. (2008). Canonical Wnt signaling regulates organ-specific assembly and differentiation of CNS vasculature. Science 322, 1247–1250. https://doi.org/10.1126/science.1164594.

Street, K., Risso, D., Fletcher, R.B., Das, D., Ngai, J., Yosef, N., Purdom, E., and Dudoit, S. (2018). Slingshot: cell lineage and pseudotime inference for single-cell transcriptomics. BMC Genomics 19, 477. https://doi.org/10.1186/s12864-018-4772-0.

Sweeney, M.D., Zhao, Z., Montagne, A., Nelson, A.R., and Zlokovic, B.V. (2019). Blood-Brain Barrier: From Physiology to Disease and Back. Physiol. Rev. 99, 21–78. https://doi.org/10.1152/physrev.00050.2017.

Travier, L., Alonso, M., Andronico, A., Hafner, L., Disson, O., Lledo, P.-M., Cauchemez, S., and Lecuit, M. (2021). Neonatal susceptibility to meningitis results from the immaturity of epithelial barriers and gut microbiota. Cell Rep. 35, 109319. https://doi.org/10.1016/j.celrep.2021.109319.

Uchida, Y., Sumiya, T., Tachikawa, M., Yamakawa, T., Murata, S., Yagi, Y., Sato, K., Stephan, A., Ito, K., Ohtsuki, S., et al. (2019). Involvement of Claudin-11 in Disruption of Blood-Brain, - Spinal Cord, and -Arachnoid Barriers in Multiple Sclerosis. Mol. Neurobiol. 56, 2039–2056. https://doi.org/10.1007/s12035-018-1207-5.

Zarbalis, K., Siegenthaler, J.A., Choe, Y., May, S.R., Peterson, A.S., and Pleasure, S.J. (2007). Cortical dysplasia and skull defects in mice with a Foxc1 allele reveal the role of meningeal differentiation in regulating cortical development. Proc. Natl. Acad. Sci. 104, 14002–14007. https://doi.org/10.1073/pnas.0702618104.

Zhou, Y., Wang, Y., Tischfield, M., Williams, J., Smallwood, P.M., Rattner, A., Taketo, M.M., and Nathans, J. (2014). Canonical WNT signaling components in vascular development and barrier formation. J. Clin. Invest. 124, 3825–3846. https://doi.org/10.1172/JCI76431.

