## Supplemental Figures 1 and 2 for "‘Formation and function of the meninges arachnoid barrier around the developing brain’"

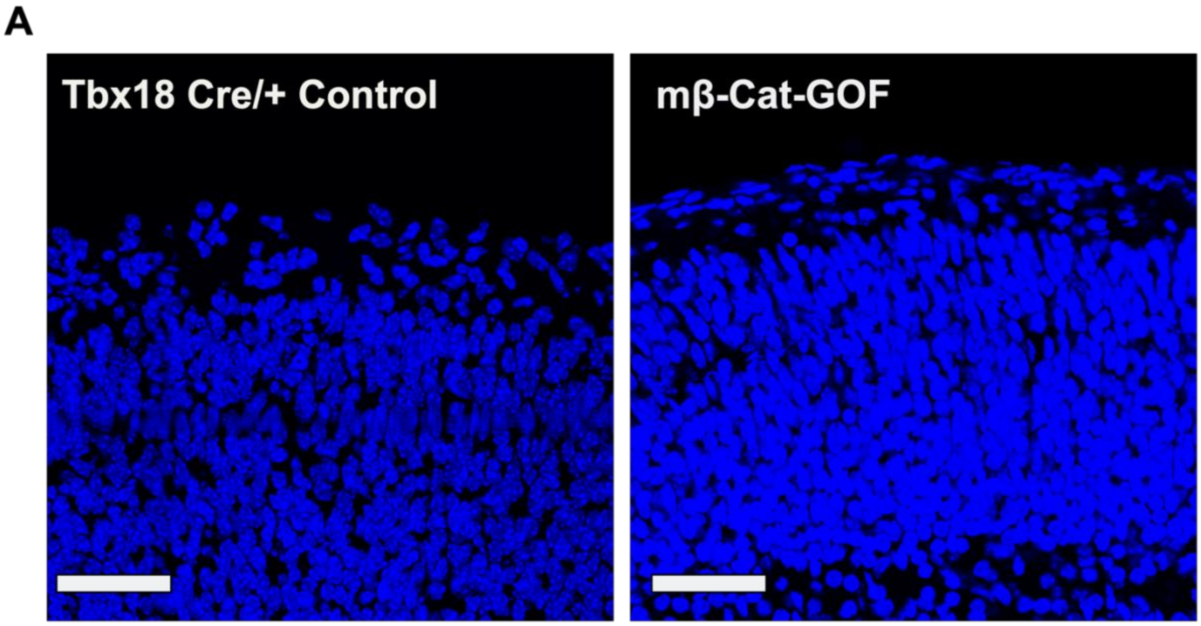


Supplemental Figure 1

DAPI staining of control and mutant brains indicating no major cortical abnormalities. Scale bar = 50 μm.

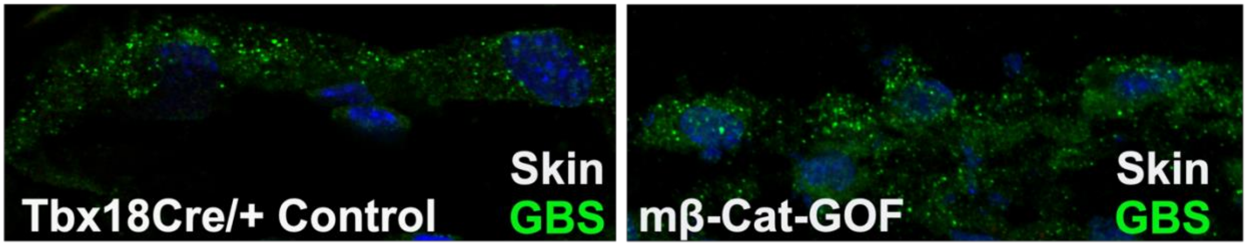


Supplemental Figure 2

Representative images from control and mutant mice 45 min after GBS inoculation (as described in Figure 2), showing comparable GBS burden in skin of mice.
